# Phosphorylation-dependent control of Arc protein by synaptic plasticity regulator TNIK

**DOI:** 10.1101/2021.04.04.438383

**Authors:** Alicyia Walczyk Mooradally, Jennifer Holborn, Karamjeet Singh, Marshall Tyler, Debasis Patnaik, Hendrik Wesseling, Nicholas J Brandon, Judith Steen, Steffen P Graether, Stephen J Haggarty, Jasmin Lalonde

## Abstract

Activity-regulated cytoskeleton-associated protein (Arc) is an immediate-early gene product that support neuroplastic changes important for cognitive function and memory formation. As a protein with homology to the retroviral Gag protein, a particular characteristic of Arc is its capacity to self-assemble into virus-like capsids that can package mRNAs and transfer those transcripts to other cells. Although a lot has been uncovered about the contributions of Arc to neuron biology and behavior, very little is known about how different functions of Arc are coordinately regulated both temporally and spatially in neurons. The answer to this question we hypothesized must involve the occurrence of different protein post-translational modifications acting to confer specificity. In this study, we used mass spectrometry and sequence prediction strategies to map novel Arc phosphorylation sites. Our approach led us to recognize serine 67 (S67) and threonine 278 (T278) as residues that can be modified by TNIK, which is a kinase abundantly expressed in neurons that shares many functional overlaps with Arc and has, along with its interacting proteins such as the NMDA receptor, been implicated as a risk factor for psychiatric disorders. Furthermore, characterization of each residue using site-directed mutagenesis to create S67 and T278 mutant variants revealed that TNIK action at those amino acids can strongly influence Arc’s subcellular distribution and self-assembly as capsids. Together, our findings reveal an unsuspected connection between Arc and TNIK. Better understanding of the interplay between these two proteins in neuronal cells could lead to new insights about apparition and progression of psychiatric disorders.

## Introduction

Neurons face constant pressure to adjust the number and strength of their connections in response to activity and other stimuli. This remarkable capacity for change involves a range of effectors whose expression and/or function can be rapidly modified in response to signals. As part of this select group of molecules, the Activity-regulated cytoskeleton-associated protein (Arc, also known as Arg3.1) is increasingly seen as a critical player because of its direct interaction with many other synaptic proteins, as well as role in multiple aspects of neuroplasticity.

Studies that examined the impact of interfering with *Arc* expression revealed the importance of this immediate-early gene for the activity of various brain systems. For instance, adult *Arc* knockout (KO) mice present lower long-term memory performance on a number of learning tasks without having problems with short-term memory (Plath *et al*. 2006), and other work also made clear that reduced *Arc* levels limit the manifestation of different types of experience-dependent neuronal change, including ocular dominance plasticity shift caused by monocular deprivation during the critical period (McCurry *et al*. 2010), the development and maintenance of binocular neurons in the visual cortex (Jenks and Shepherd 2020), and the apparition of refined spatial learning abilities at adulthood (Gao *et al*. 2018). For the most part, these different phenotypes closely align with the fact that *Arc* KO mice have deficits in long-term potentiation (LTP) (Plath *et al*. 2006; Messaoudi *et al*. 2007), long-term depression (LTD) (Waung et al. 2008; Jakkamsetti et al. 2013), and homeostatic scaling (Shepherd *et al*. 2006); as well as Arc’s known contribution to the organization and plasticity of post-synaptic terminals, which includes interaction with members of the endocytic vesicular machinery to control surface levels of 3-hydroxy-5-methyl-4-isoxazole receptors (AMPARs) (Chowdhury *et al*. 2006; Rial Verde *et al*. 2006; Dasilva *et al*. 2016), participation in actin-dependent remodeling of dendritic spines (Messaoudi et al. 2007, Peebles et al. 2010), synapse elimination in the developing cerebellum (Mikuni *et al*. 2013), and influence over the expression of AMPAR subunit *GluA1* mRNA through interaction with transcriptional regulators in the nuclear compartment (Korb *et al*. 2013). Together, these different behavioral, cellular, and molecular findings clearly illustrate the importance of Arc to brain function.

Despite more than two decades of steady discoveries about Arc expression and function, the understanding of its physicochemical and structural properties has progressed more slowly in comparison. However, a series of recent studies helped to fill major gaps concerning this topic. First, biochemical and biophysical analyses conducted with human recombinant Arc described how its modular structure, which consists of the N- and C-terminal domains divided by a flexible central hinge region, allows monomeric units to self-arrange as large soluble oligomers (Myrum *et al*. 2015). Coincidingly, Zhang and colleagues (2015) reported on their side that Arc’s C-terminal region contains two distinct subdomains, termed N-lobe and C-lobe, that share a high degree of similarity to the capsid domain of several retroviral Gag proteins, including the one seen with the human immunodeficiency virus (HIV). In addition to this observation, comparison of Arc homologs revealed conservation of the Gag domain across vertebrates—a result that supports, by the way, a prediction made a few years before about the possible retrotransposon evolutionary origin of the *Arc* gene (Campillos *et al*. 2006)— and crystal studies showed how the N-lobe region mediates intermolecular binding of Arc to the synaptic proteins Ca^2+^/calmodulin-dependent protein kinase II (CaMKII), transmembrane AMPAR regulatory protein γ2 (TARPγ2, also known as Stargazin) (Zhang *et al*. 2015), and guanylate kinase-associated protein (GKAP) (Zhang *et al*., 2015; Hallin *et al*. 2021). Further investigation within this interactive binding domain measured varying levels of binding affinities to different short peptide motifs, suggesting another layer of complexity and control in the role played by Arc as a hub protein facilitating structural rearrangement of the postsynaptic density (Hallin *et al*. 2020). Finally, these studies led the way to one of the most surprising discoveries about Arc protein which is its capacity to self-organize as virus-like capsids that can transfer genomic material, including its own mRNA transcripts, from a donor neuron to other cells (Ashley *et al*. 2018; Pastuzyn *et al*. 2018; Erlendsson *et al*. 2020). Although this specific phenomenon will require further research to understand its full significance in neuron biology, the current evidence clearly indicate that Arc should be also considered as an active participant in intercellular communication events of the nervous system.

With a role in many processes that are each mechanistically different and occurring in distinct subcellular compartments, it is not clear how Arc can be rapidly recruited to perform one specific task over another. Surely, a complete answer to this question will implicate multiple factors that guide Arc to a precise location and command its association with specialized effectors. Consistent with this point, several studies have already identified a small number of post-translational modifications (PTMs) with specific consequence on Arc. One example of this is how ubiquitination of specific lysine residues can trigger Arc proteasomal degradation (Greer *et al*. 2010; Mabb *et al*. 2014). Here, though, the influence of the ubiquitin-proteasomal system is likely more complex than initially thought as other work also revealed that the acetylation of lysine sites can inversely increase Arc protein half-life and abundance, hinting therefore at a competition between these two types of PTMs (Lalonde *et al*. 2017). In addition to these results, one study also found that SUMOylation of Arc can stimulate its association with the actin regulator drebrin A in dendritic spines to promote LTP consolidation (Nair *et al*. 2017), and another showed that preventing palmitoylation of a cluster of cysteines in Arc’s N-terminus can impact synaptic depression (Barylko *et al*. 2018). Finally, several phosphorylation events have been implicated in the control of Arc as well. These include S206 in the central hinge region targeted by Extracellular signal-regulated kinase 2 (ERK2) to seemingly control nuclear:cytoplasmic localization (Nikolaienko *et al*. 2017), multiple putative Glycogen synthase kinase-3 (GSK3α/β) sites gating degradation and effect on dendritic spines morphology (Gozdz *et al*. 2017), as well as S260 for which phosphorylation by CaMKII can prevent high-order oligomerization via interference in N-lobe and C-lobe subdomains interaction (Zhang *et al*. 2019).

These examples most certainly represent only a subset of Arc’s PTMs with many other events remaining to be found and associated with specific function. This motivated us to search for novel phosphorylation sites by combining mass spectrometry and sequence prediction strategies. This approach allowed us to identify two candidate residues, one in the N-terminal end and another at the C-terminus, that could be modified by the tumor necrosis factor receptor (Traf2) and noncatalytic region of tyrosine kinase (Nck) interacting kinase (TNIK), a member of the germinal center kinases family that is abundantly expressed in neurons and shares many functional overlaps with Arc. Using proteomics, *in vitro* assays, and overexpression experiments in mouse Neuro2a neuroblastoma cells, we collected evidence suggesting that phosphorylation of each candidate sites exert very different effects on the distribution and oligomerization of Arc. The connection between Arc and TNIK that we found provide a new direction to understand how Arc can adopt a specific role in different cellular subcompartments.

## Materials and Methods

### Cell culture and transfection

Neuro2a cells (mouse neuroblastoma cell line also known as N2a cells, RRID: CVCL_0470) were cultured in DMEM supplemented with 10% HyClone FetalClone II serum (Cytiva, Marlborough, MA, USA), penicillin (50 units/ml), and streptomycin (50 µg/ml). Cells were transfected overnight using Lipofectamine 2000 (Invitrogen, Grand Island, NY, USA) according to the manufacturer’s protocol. The Neuro2A cell line was used in previous publication by our group (Lalonde *et al*. 2017). During the experiment schedule, the cell line was subjected to 4-5 more passages.

### Antibodies and pharmacological compounds

The antibodies recognizing TNIK (1:1000 for western blotting [WB], #612210, RRID: AB_399573) and phosphoserine/threonine residues (1:1000 for WB, #612548, RRID: AB_399843) were purchased from BD Biosciences (San Jose, CA, USA). The Arc antibody (1:1000 for immunocytochemistry [ICC] and 1:2000 for WB, #156 003, RRID: AB_887694) was from Synaptic Systems (Goettingen, Germany) while the horseradish peroxidase (HRP)-conjugated FLAG (1:1000 for WB, A8592, RRID: AB_439702), FLAG M2 (1:1000 for WB, F1804, RRID: AB_262044), β-actin (1:100,000 for WB, A1978, RRID: AB_476692), and GAPDH (1:100,000 for WB, AB2302, RRID: AB_10615768) antibodies were from Sigma-Aldrich (St. Louis, MO, USA). The antibodies detecting GST (1:1000 for WB, #2625, RRID: AB_490796) and Myc-tag (1:1000 for WB, #2276, RRID: AB_331783) were acquired from Cell Signaling Technology (Beverly, MA, USA) whereas the drebrin antibody (1:500 for WB, sc-374269, RRID: AB_10990108) was from Santa Cruz Biotechnology (Santa Cruz, CA, USA). Finally, cross-absorbed HRP-conjugated secondary antibodies were from Thermo Fisher Scientific (Waltham, MA, USA).

AK-7 was purchased from Tocris Bioscience (Bristol, UK), oxamflatin from Santa Cruz Biotechnology, and KY-05009 from Sigma-Aldrich. The inactive analog G883-2176 was obtained from commercial sources (Molport, Beacon, NY, USA).

### Plasmids

The pCMV6-Arc-Myc-DDK (FLAG) mouse ORF cDNA clone (MR206218) was from OriGene Biotechnologies (Rockville, MD, USA). The pRK5 vector was a generous gift from Stephen Moss (Tufts University, Boston, MA, USA). Arc-Myc-FLAG and Arc-GST point mutants (S67A, S67D, T278A, T278D) were generated using the Q5 Site-Directed Mutagenesis Kit from New England Biolabs (Ipswich, MA, USA) according to manufacturer’s instructions. All constructs were verified by DNA sequencing.

### Western blotting

For western blotting, cells were collected by scraping in ice-cold radioimmunoprecipitation assay (RIPA) buffer (50 mM Tris-HCl [pH 8.0], 300 MM NaCl, 0.5% Igepal-630, 0.5% deoxycholic acid, 0.1% SDS, 1 MM EDTA) supplemented with a cocktail of protease inhibitors (Complete Protease Inhibitor without EDTA, Roche Applied Science, Indianapolis, IN, USA) and phosphatase inhibitors (Phosphatase Inhibitor Cocktail 3, Sigma-Aldrich). One volume of 2X Laemmli buffer (100 mM Tris-HCl [pH 6.8], 4% SDS, 0.15% bromophenol blue, 20% glycerol, 200 mM β-mercaptoethanol) was added and the extracts were boiled for 5 min. Samples were adjusted to an equal concentration after protein concentrations were determined using the BCA assay (Pierce, Thermo Fisher Scientific). Lysates were separated using sodium dodecyl sulphate-polyacrylamide gel electrophoresis (SDS-PAGE) and then transferred to a nitrocellulose membrane. Next, the membrane was blocked in TBST (Tris-buffered saline and 0.1% Tween 20) supplemented with 5% non-fat powdered milk and probed overnight at 4°C with the indicated primary antibody. Finally, after washing with TBST the membrane was incubated with the appropriate secondary antibody and visualized using enhanced chemiluminescence (ECL) reagents according to the manufacturer’s guidelines (Pierce, Thermo Fisher Scientific).

The following procedure was used to quantify western blot analyses. First, equal quantity of protein lysate was analyzed by SDS-PAGE for each biological replicate. Second, the exposure time of the film to the ECL chemiluminescence was the same for each biological replicate. Third, all the exposed films were scanned on a HP Laser Jet Pro M377dw scanner in grayscale at a resolution of 300 dpi. Fourth, the look-up table (LUT) of the scanned tiff images was inverted and the intensity of each band was individually estimated using the selection tool and the histogram function in Adobe Photoshop CC 2020 software. Finally, the intensity of each band was divided by the intensity of its respective loading control (β-actin) to provide the normalized value used for statistical analysis.

### Co-immunoprecipitation

To assess interaction between Arc and TNIK under different experimental conditions, each co-immunoprecipitation (IP) was completed with one 90% confluent 10 cm plate of Neuro2a cells overexpressing wild-type (WT) Arc-Myc-FLAG. Cells from each plate were collected and lysed in 500 µL ice-cold soft lysis buffer (20 mM Tris-HCl [pH 8.0], 150 mM NaCl, 0.2% Igepal-630, 2 mM EDTA) supplemented with a cocktail of protease inhibitors (Complete Protease Inhibitor without EDTA, Roche Applied Science, Indianapolis, IN) and phosphatase inhibitors (Phosphatase Inhibitor Cocktail 3, Sigma-Aldrich) followed by homogenization using QIAshredder spin columns (Qiagen, Hilden, Germany). Next, lysates were adjusted to a similar protein concentration after quantification with BCA assay and a fraction of each sample reserved as input material. Equal amount of the remaining lysate from each condition was used for the IP procedure. FLAG-tagged Arc was immunopurified using 50 µL of anti-FLAG M2 Magnetic Beads (Sigma-Aldrich, M8823) while endogenous TNIK was pulled-down with 10 µL of anti-TNIK antibody (BD Biosciences, #612250) and 25 µL Dynabeads Protein G (Invitrogen, Thermo Fisher Scientific, #10003D). After overnight incubation at 4°C, beads were washed with 1000 µL of soft lysis buffer thrice for 10 minutes at 4°C. For FLAG-Arc IP, the beads were resuspended in 50µL of FLAG peptide solution (1µg/µL in RIPA, Sigma-Aldrich, F3290) after the last wash to elute FLAG-tagged Arc and other proteins from beads. For TNIK IPs, beads were resuspended in 60 µL of 1X SDS-LB. Finally, samples were run according to western blotting procedure described above and 1% of protein lysate from each sample was utilized as input control.

Co-immunopreciations of Arc-Myc-FLAG (WT, S67D, and S67A) with endogenous drebrin were performed as described above. The different FLAG-tagged Arc proteins overexpressed in Neuro2a cells were immunopurified using 50 µl anti-FLAG M2 Magnetic Beads from lysates and eluted using FLAG peptide solution.

### Mass spectrometric analysis

Mass spectrometry procedure for shotgun detection of Arc phosphorylated residues was similar to analyses probing for Arc acetylation and ubiquitination modifications previously published by our group (Lalonde *et al*., 2017). In brief, WT Arc-Myc-FLAG was overexpressed in Neuro2a cells and immunopurified with anti-FLAG M2 Magnetic Beads (Sigma-Aldrich). After washes with RIPA, Arc-Myc-FLAG was eluted by incubating beads in 50 µL of RIPA buffer containing 25 µg of FLAG peptide (Sigma-Aldrich) for 2 h at 25°C with gentle agitation. Eluates from nine separate IPs were combined, concentrated by ethanol protein precipitation and separated by SDS-PAGE. After Coomassie staining, the gel band corresponding to Arc-Myc-FLAG was excised and in-gel digested using trypsin prior to mass spectrometric analysis. All LC/MS experiments were performed as detailed in Lalonde *et al*. (2017) with a Q Exactive mass spectrometer (Thermo Scientific) coupled to a micro-autosampler AS2 and a nanoflow HPLC pump (Eksigent Technologies, Dublin, CA, USA). Data for two biological replicates were processed separately and pooled.

### Recombinant protein preparation

DNA encoding WT mouse Arc was amplified from the original pCMV6-Arc-Myc-FLAG plasmid using PCR, subcloned into a pGEX-4T-3 vector between the SalI and NotI sites and transformed into the BL21 derivatives *E. coli* cells Rosetta 2 (DE3) Competent Cells (Novagen, Sigma-Aldrich). Arc S67A and T278A DNA was amplified from point-mutant pCMV6-Arc-Myc-FLAG plasmids prepared using site-directed mutagenesis and subcloned similarly. Starter bacteria cultures grown overnight at 37°C in LB supplemented with ampicillin and chloramphenicol were used to seed large volume (500 mL) cultures. Those were grown at 37°C and 300 rpm until an OD_600_ of 0.6-0.8 at which point they were induced by the addition isopropyl-β-D-thiogalactoside (IPTG) to a final concentration of 20 mM and incubated at 16°C for 16-20 h shaking (300 rpm). Subsequently, cultures were pelleted at 6,000 × *g* for 15 min at 4°C followed by resuspension of pellets in 30 mL of GST Buffer (50 mM Tris [pH 8.0], 300 mM NaCl, 10% glycerol) supplemented with 3 mM β-mercaptoethanol and 2 mM phenylmethylsulfonyl fluoride (PMSF) to limit protease activity. Resuspended cells were sonicated for 8-10 × 45 s pulses at duty load 60%, then insoluble material pelleted at 21,000 × *g* for 45 min. Supernatant was filtered with a 0.45 µm filter then left to equilibrate overnight with 1 mL of Glutathione Sepharose 4B affinity resin (GE Healthcare, Pittsburgh, PA, USA, #17-0756-01). Glutathione Sepharose 4B resin and bound protein was applied to a 10 mL disposable plastic column and washed thrice with 3 mL of wash buffer (GST Buffer + 0.1% Triton X-100). Elution of bound protein was accomplished by incubating 6 × 0.5 mL of GST Elution Buffer (50 mM Tris [pH 8.0], 300 mM NaCl, 10% glycerol, 10 mM reduced glutathione, 1 mM DTT) with the resin for 10 minutes at room temperature before elution. Finally, fractions were verified by SDS-PAGE for purified proteins and those with recombinant GST-Arc were pooled, buffer exchanged (Dialysis Buffer: 20 mM Tris [pH 7.5], 200 mM NaCl, 25% glycerol, 1 mM DTT), and concentrated using Amicon Ultra-4 Centrifugal Filer Units – 10,000 NMWL (Millipore, Sigma-Aldrich, #UFC801024) at 7,500 × *g* for 20 min.

For electron microscopy and dynamic light scattering experiments, the GST tag was cleaved using the Thrombin CleanCleave Kit (Sigma-Aldrich, SLBZ7194) according to the manufacture’s protocol after elution step. Samples were run on a HiPrep S16/60 Sephacryl S-200 HR (GE Healthcare, #17-1166-01) to separate the proteins by molecular weights and then verified by SDS-PAGE (Figure S1). Eluates containing cleaved Arc in 50 mM Tris (pH 8.0), 300 mM NaCl, and 10% glycerol were pooled and normalized.

### Kinase assay

ADP-Glo Kinase Assay reagents were from Promega (Madison, WI, USA). Reactions were performed according to manufacturer’s protocol with 1 µM full-length GST-tagged mouse Arc and 10 nM of human TNIK catalytic domain (Carna Biosciences, Japan, #07-138) in reaction buffer (50 mM Tris [pH 7.5], 5 mM MgCl_2_, 0.01% Brij-35). Reactions were initially conducted with a range of ATP concentrations (31.25, 62.5, 125, and 250 µM). For experiments with TNIK inhibitor KY-05009 and the inactive analog G883-2176, compounds were pre-incubated for 15 minutes before addition of ATP (125 *µ*M). Kinase reactions in all experiments were allowed to proceed for 60 min at room temperature and terminated by addition of ADP-Glo Reagent followed by incubating for 45 min at room temperature to deplete the remaining ATP. Next, Kinase Detection Reagent was performed for 15 min to convert ADP product to ATP and the newly synthesized ATP was measured via a luciferase/luciferin reaction with the help of luminometer plate reader.

For western blot analyses, kinase reactions performed with 400 µM ATP were terminated by adding 2X Laemmli buffer and 30 µL of each sample was run on SDS-PAGE. Membranes were then incubated for 48 h with mouse pan-phosphoserine/threonine antibody and processed as described above. Finally, to identify the specific Arc residues modified by TNIK in assay, a kinase reaction was separated by SDS-PAGE, Coomassie blue stained, and the band corresponding to GST-Arc excised and in-gel digested using trypsin followed by mass spectrometric analysis.

### Actin fractionation assay

Neuro2a cells cultured in 6-well plates overexpressing WT or point-mutants Arc-Myc-FLAG were collected with Triton X-100 lysis buffer (1% Triton X-100, 150 mM NaCl, 20 mM Tris-HCl [pH7.5]; 200 µL per well) supplemented with protease inhibitors, incubated at room temperature for 10 min and then centrifuged at 14,000 rpm for 10 min. Next, equal volume (150 μl) of supernatant representing the Triton X-100 soluble fraction with globular actin (G-actin) from each tube was collected without disrupting the pelleted material, quantified, normalized, and stored at - 80°C. In parallel, 100 μL of fresh lysis buffer was added to each tube with the non-soluble fraction and the pellet gently resuspended with pipetting followed by centrifugation at 14,000 rpm for 5 minutes. After centrifugation, the supernatant was discarded and the remaining pellet representing the insoluble filamentous actin (F-actin) fraction dissolved in 50 μL of 1X SDS-LB. Equal amount of soluble (G-actin) and insoluble (F-actin) samples were run on SDS-PAGE and the membrane probed for β-actin, FLAG, and GAPDH.

### Immunocytochemistry and actin phalloidin staining

Indirect immunofluorescence detection of antigens was carried out using Neuro2a cells cultured on glass coverslips in 6-well plate at an approximate density of 1.0 × 10^6^ cells/mL. After transfection of pRK5-Arc-GFP according to manufacturer’s protocol, cells were washed twice with phosphate-buffered saline (PBS) and fixed for 30 min at room temperature with 4% paraformaldehyde in PBS. After fixation, cells were washed twice with PBS, permeabilized with PBST (PBS and 0.25% Triton X-100) for 20 min, blocked in blocking solution (5% goat nonimmune serum and 1% bovine serum albumin in PBS) for another 30 min, and finally incubated overnight at 4°C with the primary antibody in blocking solution. The following day, coverslips were extensively washed with PBS and incubated for 2 hours at room temperature in the appropriate fluorophore-conjugated secondary antibody solution [Alexa Fluor 488- or Alexa Fluor 594-conjugated secondary antibody (Molecular Probes, Thermo Fisher Scientific) in blocking solution]. Fluorescent labelling of F-actin was performed using Alexa Fluor 594-conjugated phalloidin (Invitrogen, Thermo Fisher Scientific, A12381) according to manufacturer’s protocol. Finally, cell nuclei were counterstained with 4′,6-diamidino-2-phenylindole (DAPI), and coverslips were mounted on glass slides with ProLong Antifade reagent (Invitrogen, Molecular Probes).

Cells cultured on coverslips from three independent biological replicates were imaged with a Nikon Eclipse Ti2-E inverted microscope equipped with a motorized stage, image stitching capability, and a 60X oil immersion objective (Nikon Instruments, Melville, NY, USA). Image preparation, assembly, and analysis were performed with Nikon’s NIS-Elements, ImageJ, and Adobe Photoshop 2020. Change in contrast and evenness of the illumination was applied equally to all images presented in the study.

### Electron microscopy

For negative stain electron microscopy, 5 µL of recombinant Arc protein sample was applied to a copper 200-mesh grid for one-minute followed by removal of excess solution using filter paper. The grid was placed onto a droplet of 1% uranyl acetate for one minute, then excess wicked away. Finally, grids were air dried and imaging was performed on a Tecnai G2 F20 (FEI, Hillsboro, OR, USA). Uniformity within the grid was visually inspected before image acquisition. For measure of circumference and circularity of capsid formations, ImageJ software was used to outline manually each structure and extrapolate values according to set scale bar.

### Dynamic light scattering

Dynamic light scattering (DLS) was performed using a Malvern Zetasizer ZS (Malvern, UK). Temperature scans and size measurements were carried out at a fixed scattering angle of 173° (back scatter). Purified protein preparations were diluted to 10 μM in size exclusion buffer (150 mM NaCl, 50 mM Tris-HCl [pH 7.5]) and size measurements were made at 20°C and 30°C. Three replicates were performed for each protein, consisting of ten measurements at each temperature, with each measurement being the average of 12 runs. Data analysis was performed on intensity and volume size distribution curves and the Z-average size was calculated using Malvern DTS software. The Z-average (presented) provides a reliable measure of the mean size of the particle size distribution.

### Statistical analyses

All statistical calculations were completed with KaleidaGraph 4.5 (Synergy Software, Reading, PA, USA) or SPSS Statistics 26 (IBM, Armonk, NY). No statistical methods were employed to predetermine sample size of any of the presented experiments. Statistical analysis included normality testing, outliers test, and Levene’s test for equality of variance were completed before moving forward with parametric tests. One-way ANOVA followed by Tukey’s post hoc test for multiple comparisons were performed where indicated. A value of *p* ≤ .05 was considered statistically significant. No test for outliers was conducted and no data point was excluded. Unless mentioned otherwise, all results represent the mean ± SEM from at least three independent experiments (*n* = 3).

### Notes on Study Design

This study was exploratory and did not involve pre-registration, randomization, or blinding.

## Results

### Mapping of Arc phosphorylation by mass spectrometry

Previous studies have provided support for the phosphorylation of Arc at T175, S206, S260, and T380 in living cells (Nikolaienko *et al*. 2017; Gozdz *et al*. 2017; Zhang *et al*. 2019); however, sequence and structure-based prediction with NetPhos 3.1 (Blom *et al*. 1999) suggests the possibility that many more residues of this protein could be modified by different kinases. As an unbiased attempt to collect novel evidence of Arc serine, threonine, and tyrosine phosphorylation under biological conditions, we overexpressed mouse Arc-Myc-FLAG in Neuro2a cells, immunopurified the protein, and performed mass spectrometry. Excitingly, this effort allowed us to not only confirm phosphorylation events previously reported by other groups, like Y274 (Palacios-Moreno *et al*. 2015) and T380 (Gozdz *et al*. 2017), but also to identify 15 new candidate phosphorylation sites (Supplementary Table 1). We noted in our dataset that the Arc peptides detected 50 times or more with a specific residue phosphorylated were all located in the N- and C-termini regions of the protein (Figure 1a). Specifically, we found T7, T8, and S67 at the amino-terminus, and S366, T376, and T380 at the carboxyl-terminus, as sites that are abundantly phosphorylated in Neuro2a cells. As we were examining with attention the surrounding sequences of these six amino acids, we recognized that Arc S67 forms a recently discovered phosphorylation consensus sequence (SVGK) for TNIK (Figure 1c) (Wang *et al*. 2016)—a serine/threonine protein kinase that is highly expressed in neurons throughout the mouse brain (Burette *et al*. 2015) and considered as a key regulator of signaling pathways contributing to cognitive function (Coba *et al*. 2012). Since no evidence of a connection between Arc and TNIK had been reported to date, we consequently decided to explore the possibility of a direct biochemical interaction between these two proteins.

**Figure 1.**
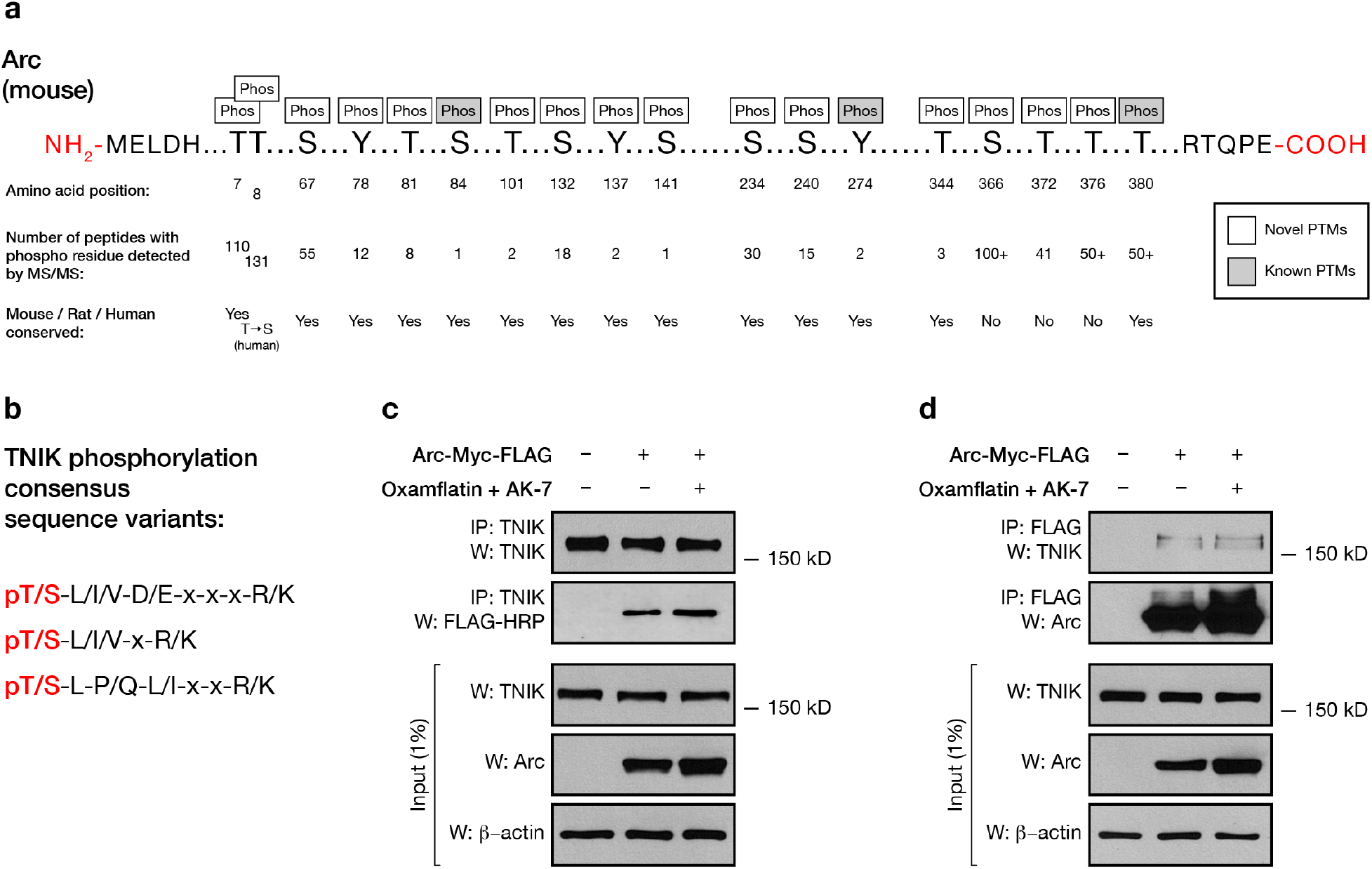
Mass spectrometry shows novel phosphorylation sites on Arc, revealing the interaction between Arc and TNIK. (a) Arc murine protein sequence displaying phosphorylated residues of Arc-Myc-FLAG immunopurified from overexpressing Neuro2a cells determined by tandem MS/MS, shaded and clear ‘Phos’ boxes represent known and novel phosphorylation events respectively. (b) TNIK phosphorylation consensus sequence variants (Wang *et al*. 2016). Residue that is phosphorylated is pictured in red. (c) TNIK coimmunoprecipitation with Myc-FLAG-Arc. The lysates of Neuro2a cells were immunoprecipitated with the anti-TNIK antibody and were analyzed by immunoblotting using anti-FLAG-HRP antibody. Input control was analyzed using TNIK and Arc, β-actin was used as a loading control. The addition of oxamflatin and AK 7 (16.67 μM) were employed to increase the abundance of endogenous Arc (Lalonde *et al*., 2017). (d) Myc-FLAG-Arc coimmunoprecipitation with endogenous TNIK. The lysates of Neuro2a cells were immunoprecipitated with the anti-FLAG antibody and were analyzed by immunoblotting using anti-TNIK and anti-Arc antibodies. Input control was analyzed using TNIK and Arc, β-actin was used as a loading control.

### Evidence of Arc as a substrate of TNIK

Direct molecular associations have been described between TNIK and several neuronal proteins, including the scaffold protein Disrupted in Schizophrenia 1 (DISC1) (Camargo *et al*. 2007; Wang *et al*. 2011), the E3 ubiquitin ligase Neuronal precursor cell expressed and developmentally downregulated protein 4-1 (Nedd4-1) (Kawabe *et al*. 2010), as well as the A-kinase anchoring protein 9 (Akap9) (Coba *et al*. 2012). To determine whether TNIK can also connect with Arc we then attempted to co-immunoprecipitate both proteins. As shown in Figure 1c, immunopurification of endogenous TNIK from Neuro2a cell lysates simultaneously pulled down overexpressed Arc-Myc-FLAG. Further, performing the same assay but in the opposite direction where FLAG-tagged Arc was purified first also isolated endogenous TNIK (Figure 1d).

These positive results motivated us to next test if this interaction between the two proteins could be extended to evidence of Arc phosphorylation by TNIK. As a starting point, we completed an exhaustive set of *in vitro* ADP-Glo kinase assays that combined bacterially expressed full-length GST-Arc and the catalytic domain of human TNIK (amino acids 1-314). Here, we measured fluorescence signals suggesting TNIK-dependent Arc phosphorylation in a manner that consistently increase with the concentration of ATP added to the assay buffer (Figure 2a). Most importantly, the measured fluorescence signal at each tested ATP concentration was significantly higher than the ones measured in the control assay reaction that combined TNIK and GST or the one that included TNIK only (Figure 2a). To confirm that the difference in fluorescence quantified between these different experimental conditions corresponded specifically to Arc phosphorylation, we then performed a western blotting analysis with ADP-Glo reaction samples and a pan-phosphoserine/threonine antibody. As seen on Figure 2b, a distinct band matching the molecular weight of GST-Arc was detected for the complete sample, but not when GST-Arc, ATP, or the TNIK catalytic domain protein was omitted from the reaction. Interestingly, a second band observed slightly below the presumed GST-Arc signal and matching the molecular weight of the TNIK catalytic fragment used the in the assay, was also detected from the complete reaction sample (Figure 2b). Since TNIK protein has four TNIK auto-phosphorylation consensus motifs, including one found between the amino acids 181-184 (TVGR), we interpreted this unexplained lower signal on the western blot as TNIK phosphorylation on itself. In order to test this hypothesis more directly, as well as further confirm that the fluorescence signal in complete ADP-Glo reaction samples corresponded to an effect of TNIK on Arc, we repeated the assay but with addition of the TNIK inhibitor KY-05009 (Figure 2c) (Kim *et al*. 2014). As expected, the presence of KY-05009 to a complete ADP-Glo reaction reduced the fluorescence signal in a dose-dependent manner when TNIK is the only protein present in the reaction or both TNIK and GST-Arc are included (Figure 2d). Of note, repeating the TNIK + GST-Arc experiment but with application of the KY-05009 inactive analog G883-2176 did not produced inhibition of the kinase reaction (Figure 2e). Finally, effect of KY-05009 on the phosphorylation of GST-Arc was also confirmed by western blot analysis probing with a pan-phosphoserine/threonine antibody (Figure 2f). Taken together, these results strongly suggest that Arc can be phosphorylated by TNIK *in vitro*.

**Figure 2.**
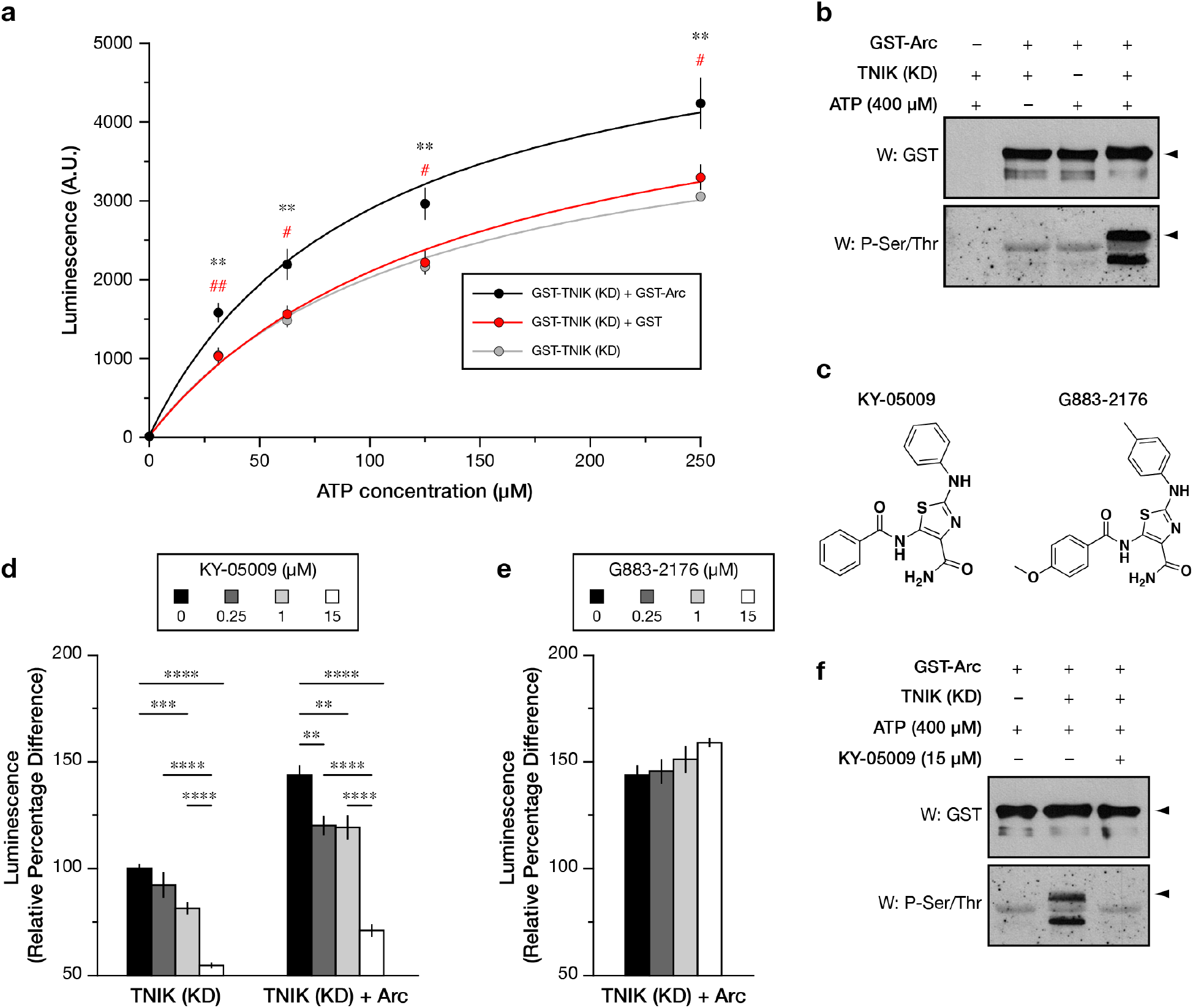
Arc was shown to be a substrate for TNIK phosphorylation using a kinase assay. (a) Arc is able to significantly increase TNIK activity with increased levels of ATP concentration in comparison to GST-TNIK alone, and control GST-TNIK + GST. One-way ANOVA revealed a significant dose–response difference between Arc and GST-TNIK (31.25μM, F_2,14_ = 10.47, *p* < 0.01; 62.5μM, F_2,14_ = 8.38, *p* < 0.01; 125 μM, F_2,14_ = 8.87, *p* < 0.01; 250 μM, F_2,14_ = 9.02, *p* < 0.01). Tukey’s HSD post hoc test, ** *p* < 0.01 in comparison to GST-TNIK (represented by the grey line), # *p* < 0.05; ## *p* < 0.01 in comparison to GST-TNIK + GST (represented by the red line). (b) Western blots showing the levels of GST to indicate Arc abundance, and phospho-serine/threonine to measure overall phosphorylation levels in Neuro2a lysate that was transfected with Arc-GST and/or TNIK and ATP (400 μM). (c) Chemical structure of TNIK inhibitor, KY-05009 (left); and inactive analog, G883-2176 (right). (d-e) KY-05009 and G883-2176 was applied in increasing concentrations (0, 0.25, 1, and 15 μM) to assess the effects on kinetic interaction between TNIK and Arc. Graphs show luminescence relative percentage difference upon addition of KY-05009 or G883-2176, respectively (±SEM). One-way ANOVA revealed a significant dose-dependent response between KY-05009 concentrations for TNIK alone (F_3,25_ = 47.92, *p* < 0.0001) as well as TNIK + GST-Arc (F_3,35_ = 37.58, *p* < 0.0001). Tukey’s HSD post hoc test, * *p* < 0.05; ** *p* < 0.01, *** *p* < 0.005; **** *p* < 0.0001. (f) Neuro2a cells were treated with KY-05009 (15 μM) and immunoblotted with anti-GST and anti-phospho-serine/threonine to analyze overall phosphorylation levels.

### TNIK modifies Arc residues at multiple sites

In order to test our hypothesis that TNIK can phosphorylate Arc at S67, we submitted GST-Arc from an ADP-Glo reaction sample to mass spectrometry analysis. As expected, Arc peptides including S67 were detected with a mass change at that location indicating phosphorylation, however, three other fragments were also detected with the amino acids S132, T278, and S366 modified in a similar fashion (Figure 3a). Two of those residues (S132 and T366) were found in our initial mass spectrometry screen (Figure 1a) but are not related to sequence arrangements currently known to be targeted by TNIK (Wang *et al*. 2016). As for T278, though, examination of the sequences immediately surrounding it revealed that this residue is, in fact, part of a TNIK phosphorylation consensus sequence (TLSR), consistent with its detection.

**Figure 3.**
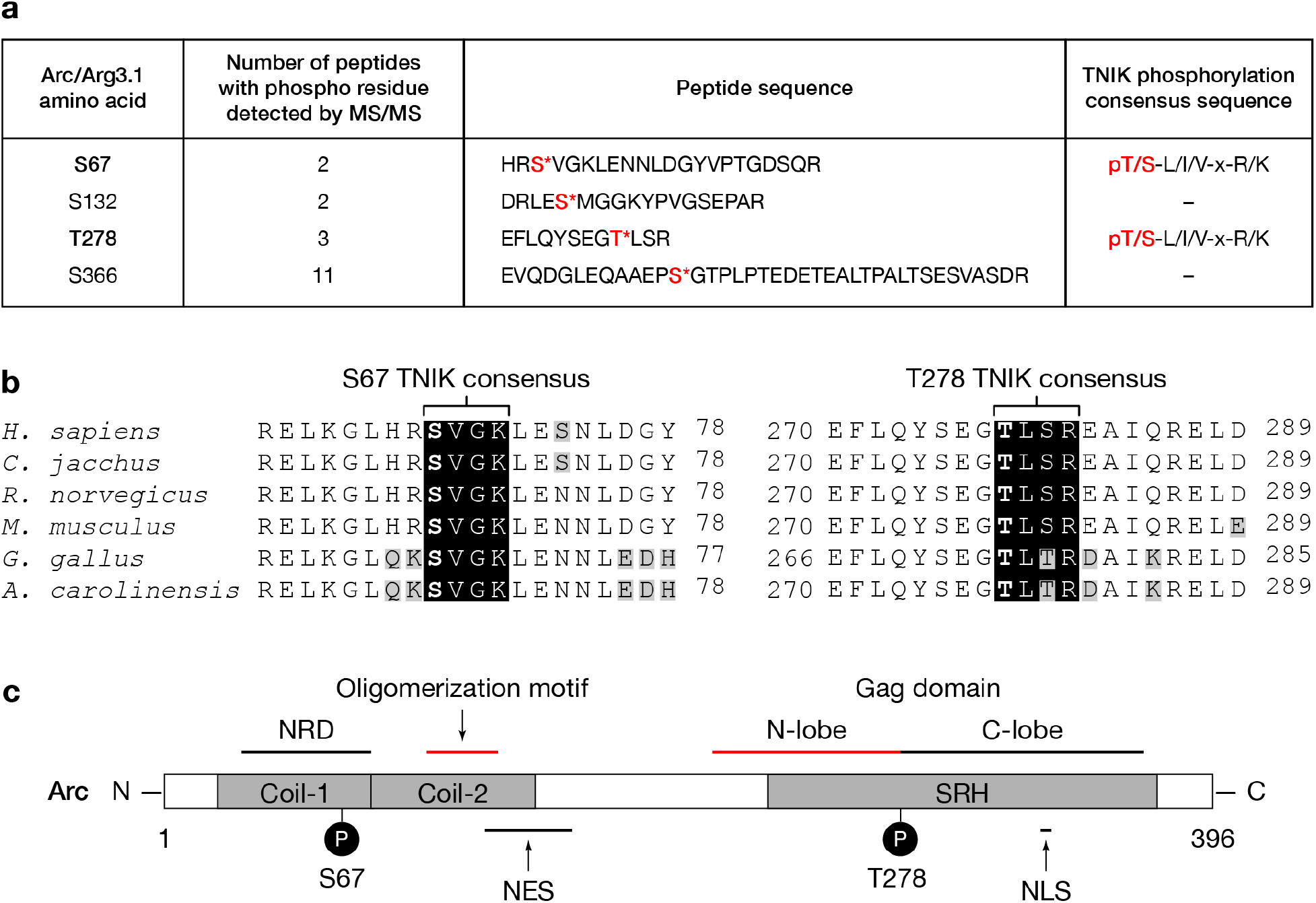
TNIK phosphorylates Arc at evolutionarily conserved sites S67 and T278. (a) Phosphorylated residues detected by MS/MS for recombinant Arc subjected to TNIK kinase assay. The phosphorylated serine (S) and threonine (T) sites in the isolated Arc tryptic peptides are presented in red with adjacent asterisk (*). (b) Alignment of Arc sequences shows that TNIK consensus sequences (black boxes) for S67 and T278 are conserved amongst a variety of species. Amino acids shaded in grey are different from consensus with other species. (c) Schematic representation of novel phosphorylation sites of interest in relation with key functional domains. NRD, nuclear retention domain (residues 29-78, Korb *et al*., 2013); NES, nuclear export signal (residues 121-154, Korb *et al*., 2013); NLS, Pat7 nuclear localization signal (residues 331-335, Korb *et al*., 2013); Coiled-coil domain (Coil-1 residues 20-77, Coil-2 residues 78-140, Eriksen *et al*., 2020); Arc oligomerization motif (residues 99-126, Eriksen *et al*., 2020); SRH, spectrin-repeat homology domain (residues 228-375); Arc Gag domain (N-lobe residues 207-277, C-lobe residues 278-370, Zhang *et al*. 2015).

Arc sequence alignment between different tetrapods show that the TNIK consensus sequences associated with S67 and T278 are both highly conserved in mammals, birds, and reptiles (Figure 3b). Most interestingly, these two residues are found on opposite sides of Arc’s central linker and within regions that have very different biophysical characteristics and function (Figure 3c). Specifically, S67 is located in the positively charged N-terminal side of the protein, within the first alpha coil (Coil-1, residues 20-77) of a predicted anti-parallel coiled-coil domain that is thought to play a role in oligomerization as well as lipid membrane binding (Hallin *et al*. 2018). In addition, work done by Korb and colleagues (2013) also provided evidence that the Arc protein segment including amino acids 29-78 could act as a nuclear retention domain (NRD). As for T278, it is located on the negatively charged C-terminal side of the protein, precisely at the transition between the N-lobe and C-lobe of the bilobar structure homologous to the retroviral Gag capsid domain (Figure 3c). Notably, the Arc N-lobe is critical to its association with postsynaptic proteins, including CaMKII and TARPγ2 (Zhang *et al*. 2016; Hallin *et al*. 2018). Based on that information, we reasoned that action of TNIK at S67 and T278 could each have very different impact on Arc biology.

### Influence of Arc phosphorylation at S67 and T278 on its distribution with F-actin

In order to gain further insights as to how phospho-modifications could influence Arc in cells, we next performed a series of experiments using overexpression of point-mutant phosphomimic and unmodifiable proteins to see how those affect its cellular distribution and oligomerization.

Evidence of interaction between Arc and the cytoskeleton protein actin include the ability of recombinant Arc to recruit F-actin from crude cellular preparations (Lyford *et al*. 1995), the inhibitory influence of Arc on the actin severing protein cofilin in the dentate gyrus (Messaoudi *et al*. 2007), as well as the fact that Arc exogenously expressed in primary hippocampal neurons localize with actin in dendritic spines and produce changes in the shape of these fine structures (Peebles et al. 2010). Based on these findings we hypothesized that phosphorylation of Arc at S67 or T278 could influence the interaction of Arc with F-actin. To test this possibility, we used Neuro2a cells overexpressing WT Arc-Myc-FLAG or a mutant version (phosphomimic or unmodifiable) of the protein for each site of interest (S67D, T278D, S67A, T278A) and performed actin co-sedimentation assays. As presented in Figures 4a and 4b, western blots for Arc with a FLAG antibody show that the five variants of the protein all had a similar expression level in the fraction with enriched G-actin (Triton X-100 soluble proteins). Most interestingly, though, probing for Arc in a similar fashion but in samples enriched for F-actin (Triton X-100 insoluble proteins) revealed that Arc unmodifiable at S67 (serine to alanine, S67A) was essentially absent from this fraction whereas the other forms of Arc (WT, S67D, T278D, and T278A) tested were similarly abundant (Figure 4a). This clear-cut result may be interpreted as the need for Arc S67 phosphorylation for its interaction with F-actin, but the performed actin co-sedimentation assay does not inform about the possibility that the Arc S67A mutant could be instead sequestered away from this major component of the cytoskeleton. To assess this possibility, we performed fluorescence immunostaining for FLAG on Neuro2a cells expressing each construct in combination with fluorescence phalloidin staining to reveal F-actin. With this approach, we first observed that WT Arc and the other mutants that co-sedimented with F-actin in our previous test (S67D, T278D, and T278A) distributed widely in transfected Neuro2a cells, including in close juxtaposition with fluorescence signals specific to phalloidin staining (Figure 4c). Interestingly, though, cells expressing Arc-Myc-FLAG S67A presented immunostaining suggesting strong accumulation of the mutant protein in the nucleus with very limited amount found in the cytoplasm (Figure 4c). Taken together, these results suggest that lack of co-sedimentation of Arc S67A with F-actin (Figure 4a) that we observed is mainly attributable to the fact that phosphorylation of this site is apparently required for trafficking of the protein out from the nucleus to the cytoplasm.

**Figure 4.**
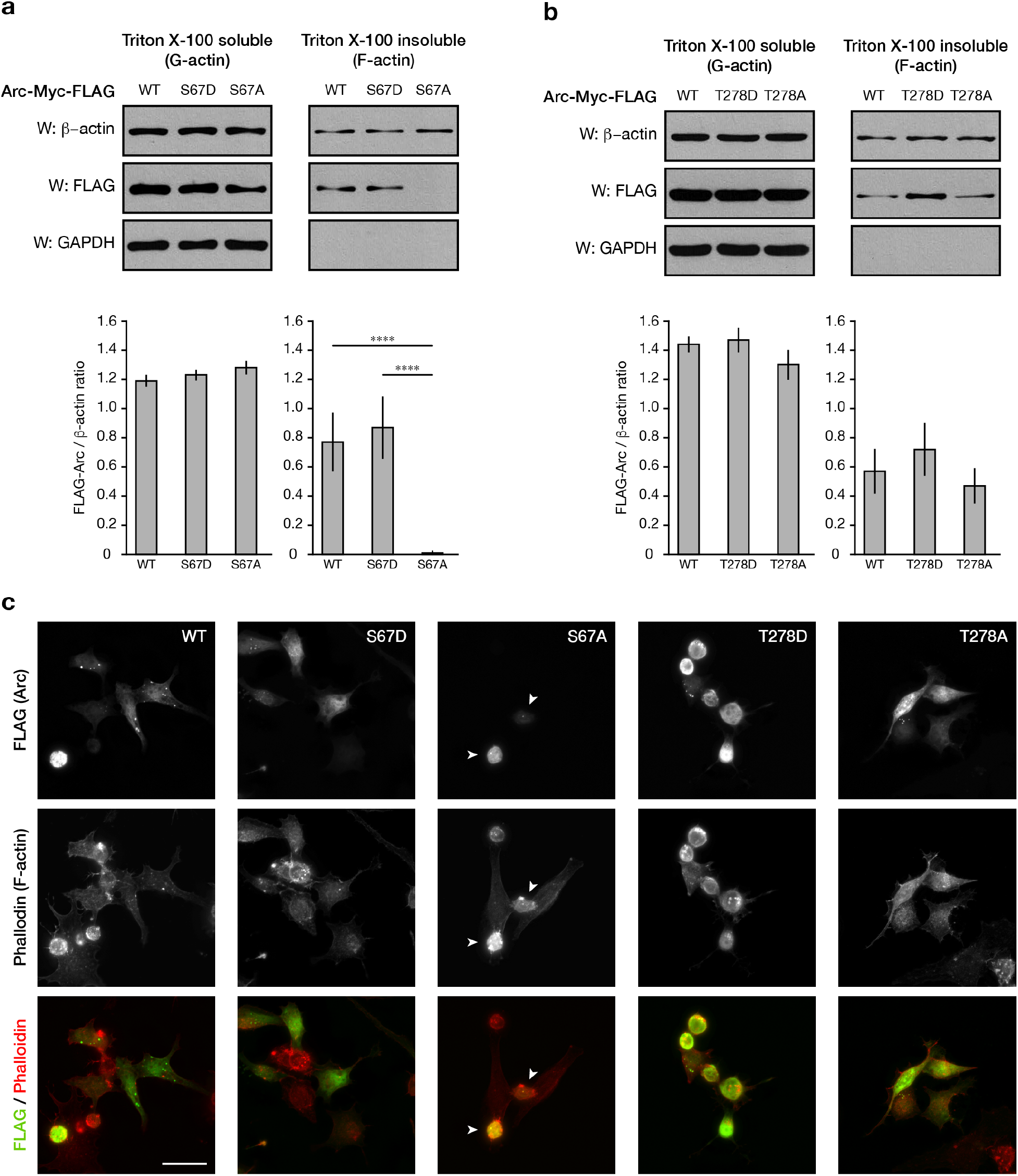
S67 influences Arc subcellular distribution. (a-b) Actin co-sedimentation assay revealed absence of unmodifiable Arc-Myc-FLAG S67A from F-actin fraction. WT, S67D phosphomimic, and both T278 variants (phosphomimic and unmodifiable) co-sedimented similarly with F-actin. Western blots show levels of β-actin, FLAG-tagged Arc, and GAPDH in Neuro2a cells that were transfected with Arc constructs and subjected to actin co-sedimentation assay. GAPDH was probed as a control and graphs show mean (±SEM) of FLAG/β-actin ratio for each condition. One-way ANOVA revealed a significant difference in distribution of Arc with F-actin for S67A with WT and S67D (F_2,11_ = 32.87, *p* < 0.0001). Tukey’s HSD post hoc test, **** *p* < 0.0001. (c) Neuro2a cells were transfected with constructs to express Arc-Myc-FLAG (WT or the different point-mutants). Representative images of cells fixed, immunostained with an antibody recognizing FLAG to detect exogenously expressed Arc (top row) and incubated with Alexa Fluor 594-conjugated phalloidin to reveal F-actin distribution (middle row) The merged microscopy captures (FLAG green fluorophore, phalloidin red fluorophore) are presented in the bottom row. Arc-Myc-FLAG S67A (arrowheads), but not the other tested variants, appeared to be only found the nuclei of transfected cells. Scale bar 250 µm.

### Modification of Arc S67 attenuates its interaction with Drebrin

Drebrin contributes to dendritic spine growth and plasticity by favoring formation of stable actin filaments (Koganezawa *et al*. 2017). Of note, Nair and colleagues (2017) have shown direct binding of Arc with drebrin A during LTP consolidation, and other research has also uncovered a role for this F-actin binding protein in trafficking of the CaMKII beta subunit (Yamazaki *et al*. 2018)—a protein that, interestingly, is also known to specifically sequester Arc at inactive synapses (Okuno *et al*. 2012). Considering those molecular connections, we then tested whether endogenous drebrin interaction with overexpressed Arc-Myc-FLAG would be affected when S67 is mutated as an aspartic acid (phosphomimic) or alanine (unmodifiable) mutant. Consistent with our previous actin fractionation and fluorescence immunostaining experiments, co-immunoprecipitation of drebrin from Neuro2a cells with exogenously expressed Arc-Myc-FLAG was strongly reduced with the unmodifiable S67A Arc variant (Figure 5).

**Figure 5.**
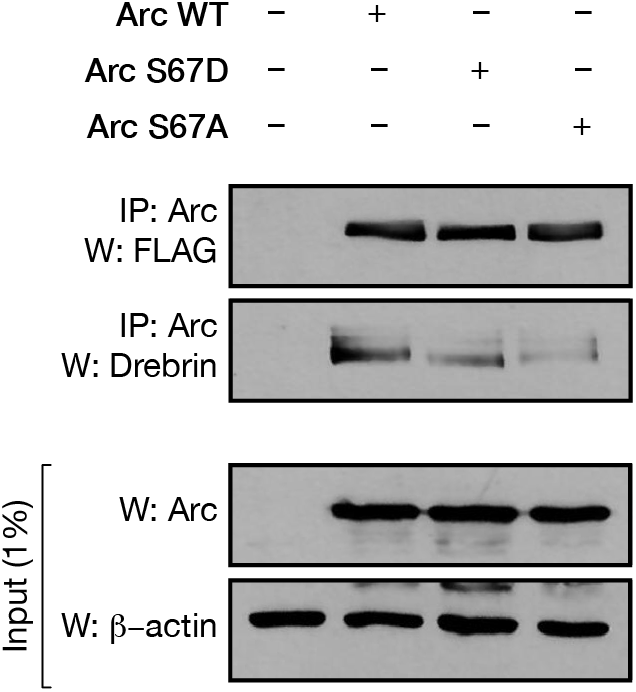
Modification of S67 influences interaction with F-actin binding protein Drebrin. Myc-FLAG-Arc coimmunoprecipitation with endogenous drebrin. The lysates of Neuro2a cells were immunoprecipitated with the anti-FLAG antibody and were analyzed by immunoblotting using anti-Arc and anti-drebrin antibodies. Input samples were analyzed using drebrin and Arc antibodies, and β-actin was used as a loading control.

### Phosphomimics of Arc S67 and T278 affect virus-like capsid formation differently

Arc can self-assemble as particles resembling retroviral Gag capsids (Ashley *et al*. 2018; Pastuzyn *et al*. 2018; Erlendsson *et al*. 2020). For mammalian Arc, capsid formation requires the second alpha-helix (Coil-2, amino acids 78-140) of the N-terminal coiled-coil assembly (Eriksen *et al*., 2020) which allows association between the N-terminal region of one Arc monomer with the C-terminus of another (Byers *et al*. 2015; Myrum *et al*. 2015; Hallin *et al*. 2018) Since S67 is found just before the Coil-2 amino acid stretch mediating self-association, and that T278 is centrally positioned in the C-terminus segment that binds with the N-terminal region, change in charge caused by phosphorylation of either residue could have profound impact on how Arc monomers organize as capsid-like structures. To examine this possibility, we performed negative-stain transmission electron microscopy (EM) with recombinant Arc and noticed that capsids produced with WT and S67D variants have round, regular sized appearance while T278D appeared as large, irregular shaped aggregates (Figure 6a-c). Supporting these observations, we quantified that the circumference of WT and S67D Arc capsids had comparable average sizes of 65.04 nm and 68.73 nm, respectively, whereas T278D were significantly larger with an average span of 184.5 nm (Figure 6d). For circularity, we used built-in ImageJ calculation (scale of 0-1 where 1 is perfectly circular and 0 is a line) and found that Arc S67D capsids presented a significantly higher circularity score (0.91) than both WT (0.85) and T278D (0.65) examples, with the last two conditions also significantly different from each other (Figure 6e). Finally, to corroborate the effect of the S67D and T278D point-mutations on homogeneity and oligomerization of Arc we performed dynamic light scattering (DLS) analysis. With this approach, we measured a size distribution of 27.5 ± 5.0nm (35%) and 307.7 ± 173.0 nm (65%) at 20°C, and 50.2 ± 32.7 nm (26%) and 333.00± 68.4 nm (74%) at 30°C for WT Arc; 38.9 ± 1.5 nm (48%) and 274.8 ± 119.6 nm (52%) at 20°C, and 38.9 ± 1.5 nm (52%) and 294.6 ± 105.2 nm (48%) at 30°C for Arc S67D; and 25.4 ± 7.9 nm (13%) and 423.7 ± 47.8 nm (87%) at 20 °C and 376.3 ± 29.9 nm (100%) at 30°C for Arc T278D (Figure 7f). In sum, those results are consistent with a previous DLS experiment for WT Arc (Myrum *et al*., 2015), and confirm the strong tendency of T278D Arc to organize as large aggregates.

**Figure 6.**
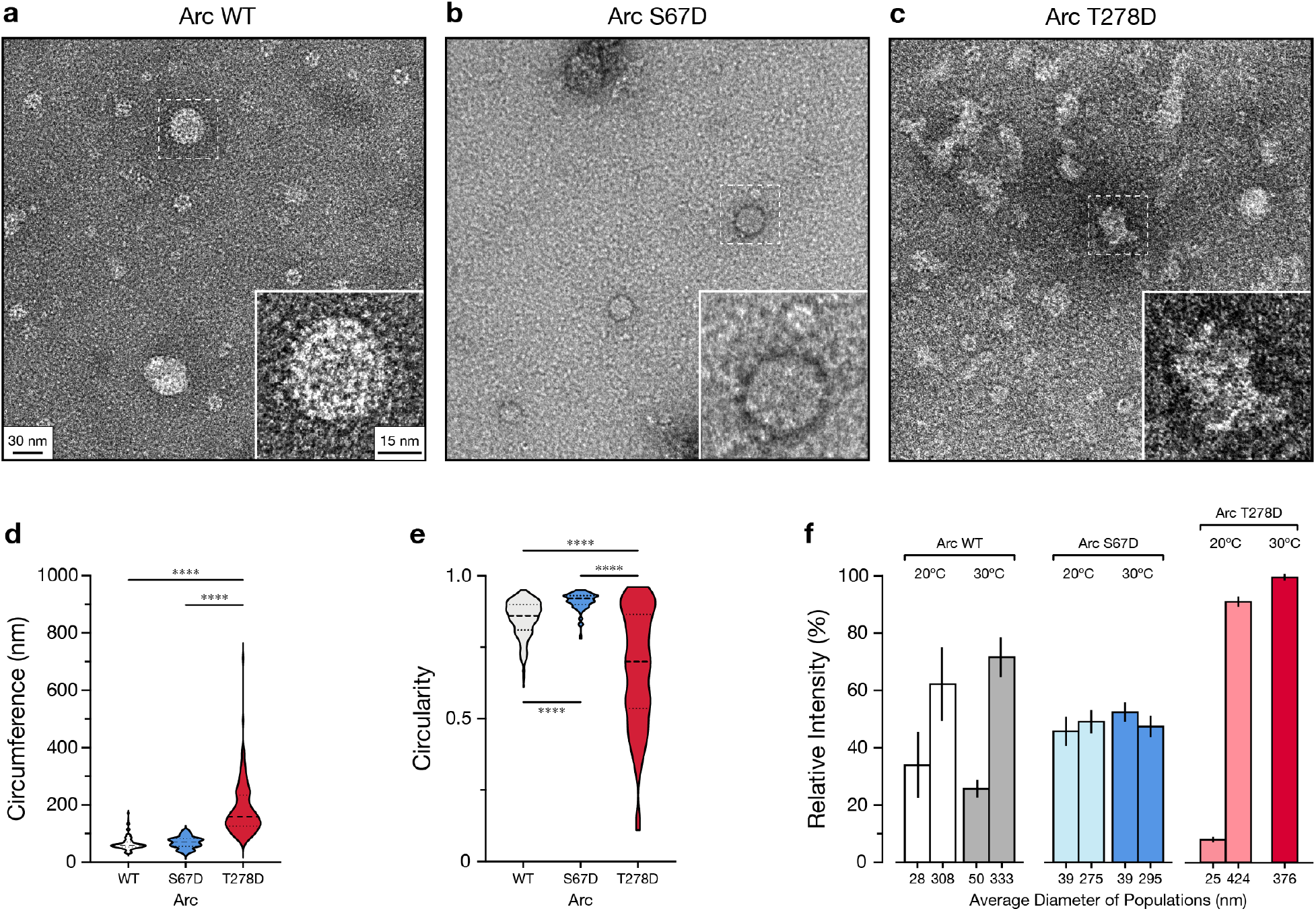
T278 plays a role in capsid dynamics. (a-c) Negative stain EM images (48,000X magnification) of capsid formations prepared with WT (a), S67D (b), and T278D recombinant Arc. (d-e) Violin plots comparing circumference (d) and capsid circularity (e) for each Arc variant. The dashed line represents the median while the dotted lines represent the two quartile lines. One-way ANOVAs revealed significant main effects (circumference, F_2, 535_ = 294.36, *p* < 0.0001; circularity, F_2, 535_ = 150.8, *p* < 0.0001). Tukey’s HSD post hoc test, *** *p* < 0.005; **** *p* < 0.0001. (f) DLS-derived weighted size distribution for each Arc variant at two temperatures (20°C and 30°C) represented as bar graph wit x-axis indicating average diameter of measured subpopulations and error bars representing SEM.

**Figure 7.**
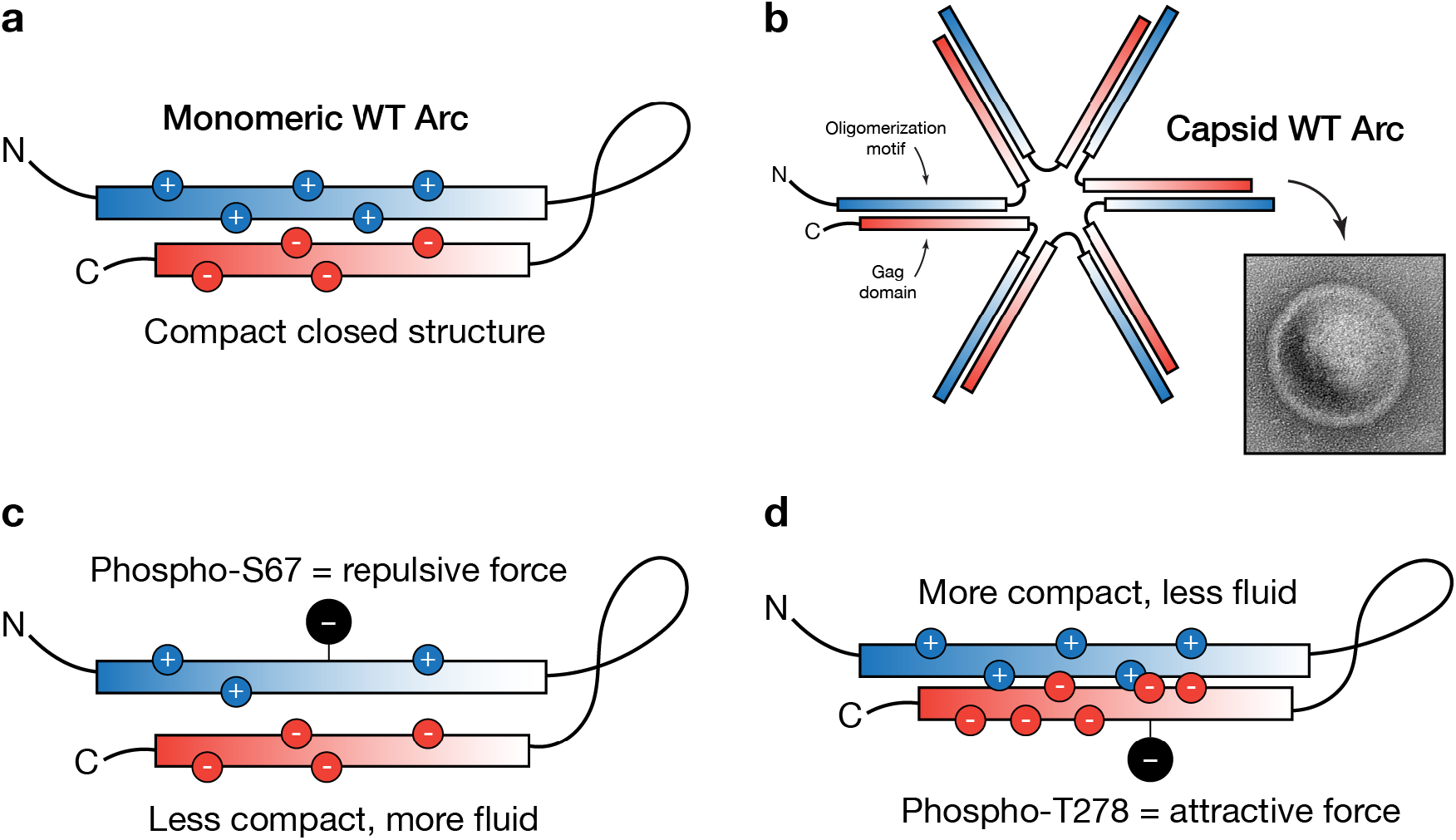
Arc monomeric and oligomeric organization. (a) Monomeric Arc form a compact closed structure where N-terminal oligomerization motif binds to C-terminus Gag domain. (b) Arc monomers can form virus-like capsids (EM image capture shown) by establishing domain swapping interactions between the N-terminal of one unit and the C-terminus of another. (c-d) Predicted influence of S67 (c) and T278 (d) phosphorylation on the contact affinity of an Arc monomer.

## Discussion

As a hub protein expressed in different parts of neuronal cells, very little is known about what guides Arc towards specific interactions and processes both spatially and temporally. Postulating that PTMs could play a pivotal role in providing specificity, we probed exogenous Arc-Myc-FLAG immunopurified from Neuro2a cells with mass spectrometry to identify residues modified by phosphorylation. While considering the various candidates that we had detected with this approach, we recognized S67 and T278 as possible targets for TNIK—a kinase that had not been directly investigated in relation to Arc even though both proteins share tantalizing similarities concerning their expression and contribution to neuron biology. Using *in vitro* kinase assays and proteomics, we confirmed that TNIK can indeed phosphorylate Arc at S67 and T278. Furthermore, we uncovered that both amino acids strongly influence, each in their own way, Arc’s subcellular distribution and/or oligomerization as virus-like capsids. Above all, our study provides evidence for a potential multifaceted interplay between Arc and TNIK. Better understanding of their connection in neuronal cells will assuredly provide valuable new insights about synaptic plasticity, the molecular underpinnings of cognition, and how disruption of their interaction could be an important factor in the apparition and progression of certain brain disorders.

### Neuronal TNIK

TNIK is highly expressed in the mammalian brain where it is not only enriched in postsynaptic densities and synaptosomal fractions (Jordan *et al*. 2004; Peng *et al*. 2004; Trinidad *et al*. 2008), but also found in neuronal nuclei where it is an active regulator of protein complex formation (Coba *et al*. 2012). Notably, Burette and colleagues (2015) have provided convincing microscopy evidence confirming accumulation of TNIK in postsynaptic densities throughout the adult mouse brain, which aligns with results showing that TNIK KO mice have significant learning deficits and altered synaptic function (Coba *et al*. 2012). At the molecular level, TNIK was found to interact with Disc1 to regulate key synaptic proteins like glutamate receptors and postsynaptic density protein 95 (PSD-95) (Camargo *et al*. 2007; Wang *et al*. 2011). Finally, it is also known that TNIK is an effector of the GTPase Rap2 through which it regulates dendrite patterning and synapse formation (Taira *et al*. 2004; Hussain *et al*. 2010). Together, those results support the exciting possibility that neuronal TNIK could act on different Arc residues in a manner that is specific to each subcellular compartment where both proteins overlap.

### Arc S67 phosphorylation and trafficking to nuclear compartment

Our study focused on Arc S67 and T278 as direct targets of TNIK activity. Taken together, our findings show that phosphorylation of each site produces very unique effects in Neuro2a cells. Starting with S67, which is found within the first alpha coil of Arc’s coiled-coil assembly (Figure 3c), we observed a drastic loss of co-sedimentation of Arc with F-actin when this residue is modified to alanine to produce an unmodifiable point-mutant variant. Importantly, immunostaining and co-immunoprecipitation experiments suggest that this result is not attributable to loss of Arc interacting with F-actin, but rather to the sequestration of the mutant protein in the nuclear compartment. A possible interpretation of why Arc S67A strongly accumulates in the nucleus is provided by the discovery of a NRD in Coil-1 that includes residue S67 (Korb *et al*. 2013). Precisely, although Arc is small enough to diffuse into the nucleus, its import is apparently regulated by a Pat7 nuclear localization signal at amino acids 331-335 (Figure 3c). Once inside the nuclear compartment, evidence suggest that Arc is retained there by the NRD which limits the activity of an adjacent nuclear export signal (NES) found at residues 121-154 by favoring interactions with other molecular components (Korb *et al*. 2013). Interestingly, our finding of a quasi-complete sequestration of unmodifiable S67A Arc suggest that phosphorylation at this specific site is required for the release of Arc from its nuclear interactors to then allow export. Known nuclear molecular interactions of Arc include the formation of a complex composed of the β-spectrin isoform βSpIVΣ5, promyelocytic leukemia (PML) bodies, and acetyltransferase Tip60 that organize to increase acetylation of histone 4 at lysine 12 (Bloomer *et al*. 2007; Wee *et al*. 2014). Whether TNIK could intervene as a negative regulator of this process by phosphorylating Arc at S67 to stimulate its nuclear export is an interesting possibility worth future investigation.

### Influence of S67 and T278 on Arc oligomeric status

In addition to the characterization of its subcellular distribution in relation to F-actin, we also analyzed how Arc recombinant protein can self-assemble as virus-like capsids using EM imaging. With this approach, we found an effect for S67 where mimicking phosphorylation of that site (S67D) made the capsids more circular, but not different in terms of average circumference, when compared to those obtained with WT Arc. Our most striking result, however, was collected with the T278D phosphomimic variant where oligomeric structures were significantly bigger and irregular than those prepared with S67D or WT (Figure 6d-e). This result can be explained by the biophysical properties of Arc and how those are possibly affected differently by S67 and T278 phosphorylation. As illustrated in Figure 7a, monomeric full-length Arc is a compact, closed structure in which the oppositely charged N-terminal domain (positive) and C-terminus region (negative) are juxtaposed, and the flexible linker between them is compacted (Hallin *et al*. 2018). When Arc molecules assemble as capsids, the same principle applies but in a domain swapping manner where the N-terminal and C-terminus of distinct monomers connect together (Figure 7b). Considering this information, it is important to recognize that phosphorylation of S67 will result in a decrease of the N-terminal positive charge (Figure 7c), making presumably molecular exchange more fluid, while phosphorylation of T278 will inversely result in stronger affinity due to increased net negative charge of the C-terminus (Figure 7d). Since Arc oligomerization involves domain swapping assembly between the N-terminal and C-terminus amongst monomers, it is then not surprising that mimicking phosphorylation at S67 and T278 separately produce very different effects. For S67, exchange between individual Arc molecules is expected to be more evenly distributed because of the reduced interaction strength between binding units. On the other hand, phosphorylation of T278 will cause tighter binding making dissociation between Arc molecules more difficult once formed. In other words, the presence of larger aggregate-like assemblies that we measured with the Arc T278D variant should be attributed to greater contact affinity with reduced likelihood of dissociation. Finally, the fact that no capsid-like structures with the expected average size (about 32 nm according to Pastuzyn *et al*., 2018) were measured for Arc T278D at a temperature (30°C) in our DLS data further support this interpretation.

### Implications of TNIK-dependent Arc phosphorylation for neuron biology and brain disorders

Our study supports the need for a systematic search and evaluation of TNIK-dependent Arc phosphorylation events in neuronal cells. As already hypothesized above, one possible function of TNIK in the nucleus could be to promote Arc export to the cytoplasm by limiting the influence of its NRD via S67 phosphorylation. In dendrites and postsynaptic structures, our accumulated knowledge about Arc and TNIK suggest that a direct relationship between those two proteins could unfold in many ways with important ramifications to neuroplasticity, cognition, and behavior. A scenario involving modification of T278 would be to alter Arc’s association with synaptic effectors that have been recognized to bind its C-terminus Gag domain. Intermolecular interactions deserving close attention include TARPγ2 (Zhang *et al*. 2016), a transmembrane protein that connects with AMPARs and mediate critical aspects of their trafficking and gating properties (Jackson and Nicoll 2011), as well as Psd-95 that can recruit Arc in an activity-dependent manner to postsynaptic densities to create supercomplexes with neurotransmitter receptors and auxiliary proteins (Fernandez *et al*. 2017). In line with the idea of TNIK targeting T278 to alter synaptic interactions, it is interesting to highlight that Arc association with the GluN2A and GluN2B subunits of N-methyl-D-aspartate-type glutamate receptors (NMDARs) has been recently found to favor stabilization of the monomeric state and prevent the formation of higher order oligomeric structures (Nielsen *et al*. 2019). Hence, phosphorylation of T278 in this case would be expected to diminish monomeric Arc association with NMDARs to promote instead formation of oligomeric structures for intercellular communication. Finally, it is critical to consider our speculations about TNIK-dependent Arc phosphorylation in light of the fact that the occurrence of Arc T278 phosphorylation was previously reported by two separate studies using a mass spectrometry approach, including global characterization of murine synaptosomes for O-GlcNAcylation and phosphorylation (Trinidad *et al*. 2012), and more recently with endogenous Arc immunopurified from adult WT mouse forebrain (Zhang *et al*. 2019) Although Arc T278 phosphorylation from brain and cultured neurons has not been validated yet with a specific phospho-Arc Thr278 antibody, neither its origin related to the activity of a specific kinase, these efforts certainly provide a degree of assurance that this site fulfill a functional role *in vivo*.

One final point important to highlight is that Zhang and colleagues (2019) also used DLS in their work to evaluate the oligomerization of Arc T278D recombinant protein and report the formation of tetramers at 20°C and high-order oligomers at 30°C, but only of smaller size than WT Arc. This is, interestingly, different from our result of only large aggregate-like formation with Arc T278D at 30°C—an observation that we are supporting with EM evidence (Figure 6). We believe that this discrepancy between the two studies could be attributable to technical differences in the preparation of recombinant GST-Arc, tag cleavage, and/or composition of buffers used in the DLS experiment. Specifically, Zhang and colleagues (2019) included reducing (DTT) and chelating (EDTA) agents in their buffer for DLS experiment, which we did not do to be consistent with our EM buffer condition. Furthermore, we noticed when optimizing our DLS protocol that filtering the recombinant protein solutions with a 0.22 µM syringe filter before analysis almost completely eliminated the measured T278D large population species at 30°C, which led us to omit this step for all conditions in our final analysis. Hence, performing such manipulation could have influenced the overall presence of protein aggregates within preparations and influenced end results.

In summary, better understanding TNIK-dependent Arc phosphorylation in neurons could offer valuable new insights about brain disorders, in particular those like schizophrenia for which accumulating evidence suggest that aberrant synaptic Arc molecular interactions are contributing to disease apparition and progression (Managò and Papaleo 2017). In line with this possibility, uncontrolled TNIK activity, which is thought to occur as a result of disruptive mutations to the psychiatric disease risk factor Disc1 (Wang *et al*. 2011), could then directly extend to major functional changes on downstream neuronal substrates like Arc. Given the dynamic nature of Arc-mediated neuroplasticity, future studies seeking to probe the potential role of TNIK in regulating Arc oligomerization, self-assembly as virus-like capsids, and interactions with synaptic proteins like the NMDA receptor, will benefit from using newly developed, selective inhibitors and other modulators of TNIK kinase activity that are under development with the long-term goal of validating potential novel therapeutic targets for the neuropsychiatric disorders (Read *et al*. 2019).

## Acknowledgements and conflict of interest disclosure

This project was supported by the Natural Sciences and Engineering Research Council of Canada (NSERC grant 401389 to J.L.), the Canadian Foundation for Innovation (CFI grant 037755 to J.L.), AstraZeneca postdoctoral program (J.L./S.J.H.), NIMH/NIH R01MH095088 (S.J.H.) and the Stuart & Suzanne Steele MGH Research Scholars Program (S.J.H.). A.W.M was supported by the University of Guelph (Graduate Tuition Scholarship). In memoriam of Robert (Bob) Harris (Advanced Analysis Centre, University of Guelph) who provided generous assistance with electron microscopy.

S.J.H. was or is a member of the scientific advisory board of Rodin Therapeutics, Psy Therapeutics, Frequency Therapeutics, Vesigen Therapeutics and Souvien Therapeutics, none of whom were involved in this study. S.J.H. has also received speaking or consulting fees from Amgen, AstraZeneca, Biogen, Merck, Regenacy Pharmaceuticals, Syros Pharmaceuticals, as well as sponsored research or gift funding from AstraZeneca, JW Pharmaceuticals, Juvenescence, and Vesigen unrelated to the content of this manuscript. N.J.B. was an employee and shareholders of AstraZeneca during the time the experiments described herein were conducted. The other authors declare no conflict of interest.

The authors declare this study was not pre-registered.

## Institutional approval

Institutional approval was not required for this study; the experiments were approved at the national level.

## Abbreviations

AMPAR: α-amino-3-hydroxy-5-methyl-4-isoxazolepropionic acid receptor
Arc: activity-regulated cytoskeleton-associated protein
BSA: bovine serum albumin
CaMKII: calcium/calmodulin-dependent protein kinase II
DAPI: 4′,6-diamidino-2-phenylindole
DISC1: disrupted in schizophrenia 1
DLS: dynamic light scattering
DMEM: Dulbecco’s modified eagle’s medium
DTT: dithiothreitol
ECL: enhanced chemiluminescence
EDTA: ethylenediaminetetraacetic acid
FPLC: fast protein liquid chromatography
GAPDH: glyceraldehyde 3-phosphate dehydrogenase
HIV: human immunodeficiency virus
HPLC: high performance liquid chromatography
HRP: horseradish peroxidase
ICC: immunocytochemistry
IP: immunoprecipitation
IPTG: isopropyl β-D-1-thiogalactopyranoside
KO: knockout
LB: Luria–Bertani medium
LC/MS: liquid chromatography–mass spectrometry
LTD: long-term depression
LTP: long-term potentiation
mGLUR: metabotropic glutamate receptor
NRD: nuclear retention domain
PCR: polymerase chain reaction
PMSF: phenylmethylsulfonyl fluoride
PSD-95: postsynaptic density protein 95
PTMs: post-translational modifications
RIPA: radioimmunoprecipitation assay
SDS-PAGE: sodium dodecyl sulphate-polyacrylamide gel electrophoresis
TARP*γ*2: transmembrane AMPAR regulatory protein *γ*2
TNIK: the tumor necrosis factor receptor (Traf2) and noncatalytic region of tyrosine kinase (Nck) interacting kinase
WB: western blot
WT: wildtype

---

**Figure S1.**
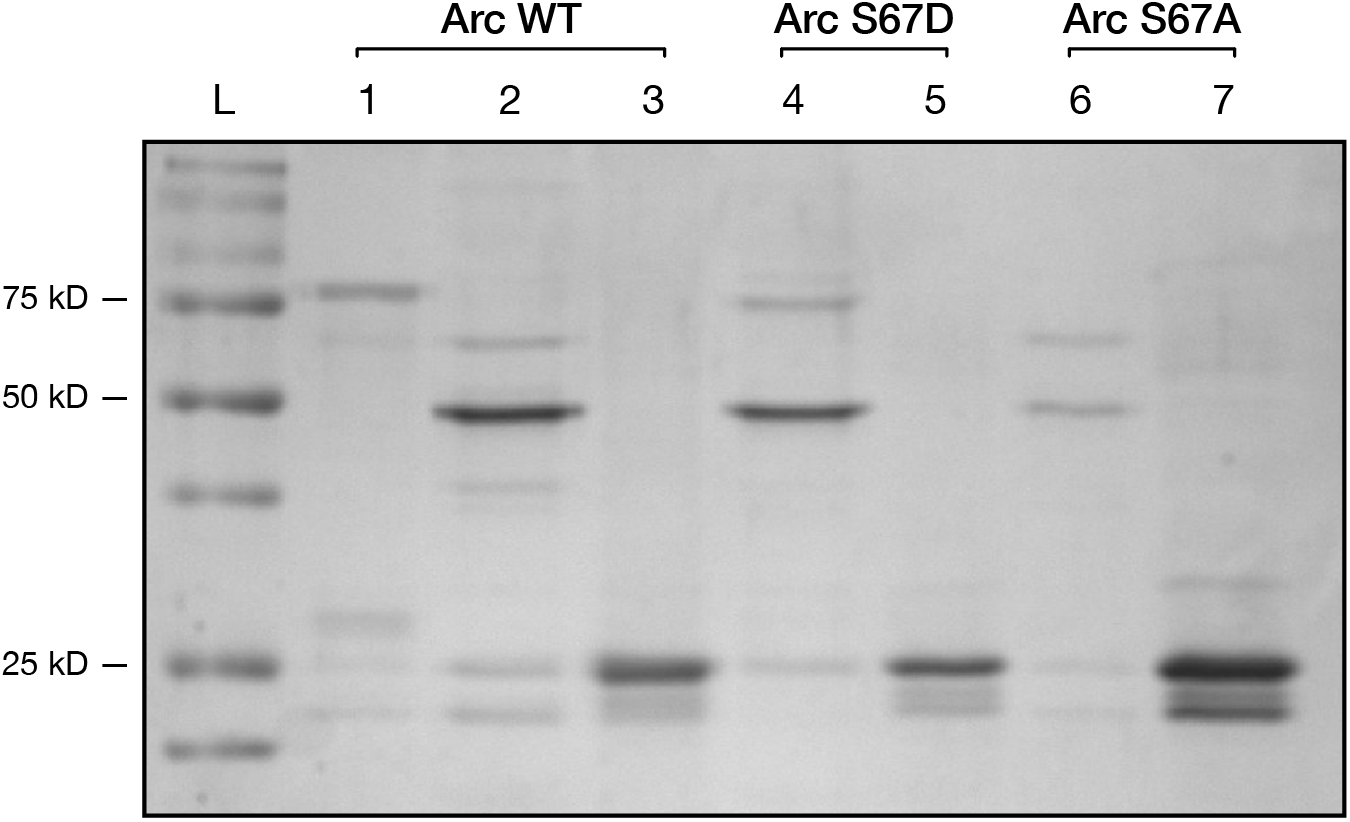
Coomassie gel showing purification of cleaved Arc variants. Lane 1) uncleaved WT Arc-GST loaded onto SEC; 2) cleaved WT Arc concentrated fractions; 3) cleaved GST from WT Arc; 4) cleaved S67D Arc concentrated fractions; 5) cleaved GST from S67D Arc; 6) cleaved T278D Arc concentrated fractions; 7) cleaved GST from T278D Arc.

**Supplementary Table 1.**
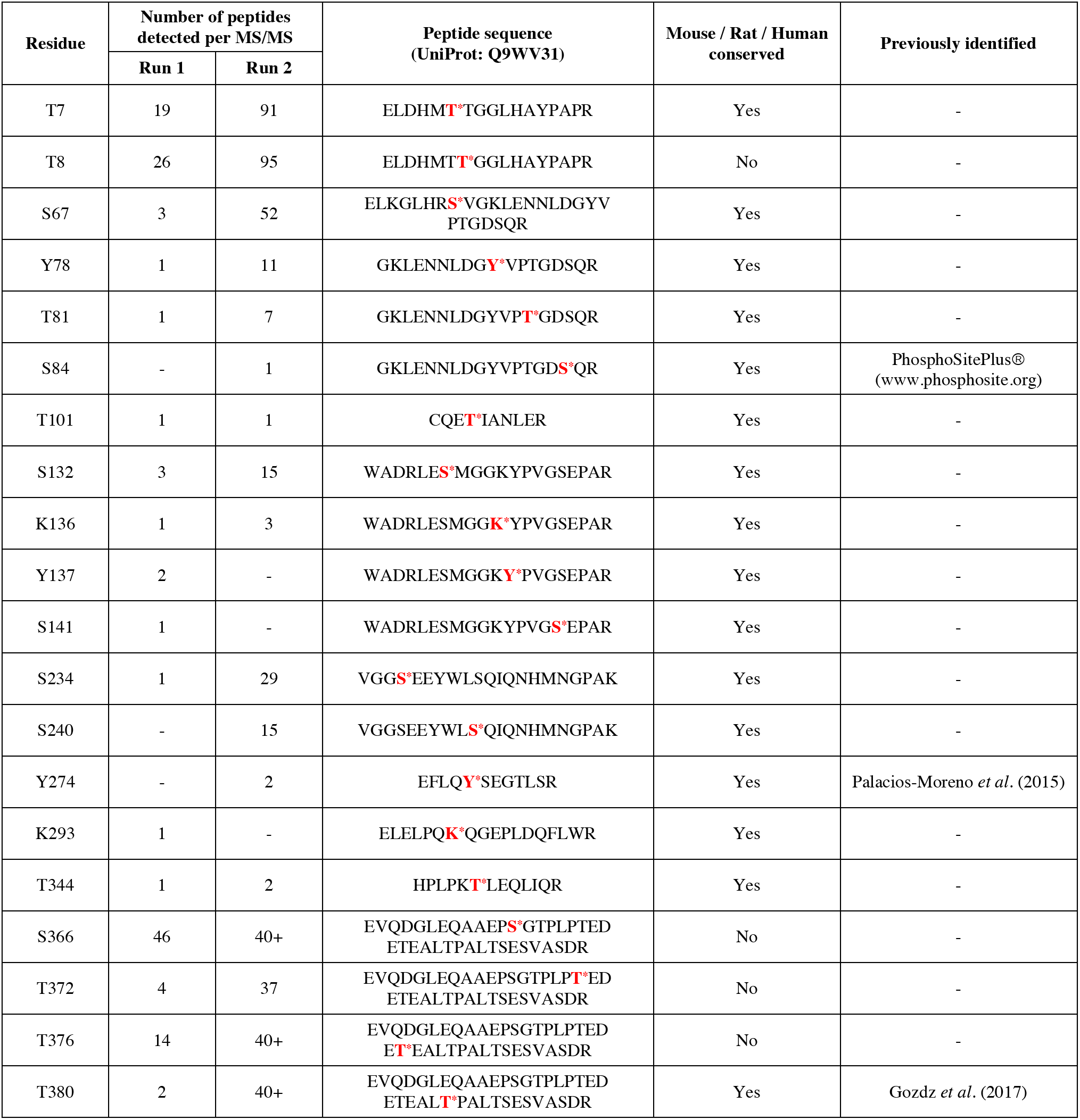
Phosphorylation peptide information pertaining to results obtained from MS/MS from Neuro2a cells overexpressing WT Arc-Myc-FLAG. The phosphorylated serine (S) / threonine (T) / tyrosine (Y) residues of the isolated Arc tryptic peptides are presented in red with asterisk (*). The total number of peptides detected with a phosphorylation modification is presented separately for each independent biological replicate.

